# Comprehensive analysis of nonsense-mediated mRNA decay targets and activity in cardiomyocytes

**DOI:** 10.64898/2026.01.12.699048

**Authors:** Mohsen Naghizadeh, Ahmed Alameldeen, Rebecca Kistler, Doris Lindner, Verena Kamuf-Schenk, Mirko Völkers, Johanna Schott, Timon Seeger, Georg Stoecklin

**Affiliations:** Division of Biochemistry, Mannheim Institute for Innate Immunoscience (MI3), Medical Faculty Mannheim, Heidelberg University, 68167 Mannheim, Germany; Center for Molecular Biology of Heidelberg University (ZMBH), German Cancer Research Center (DKFZ)-ZMBH Alliance, 69120 Heidelberg, Germany; Department of Cardiology, Angiology and Pneumology, University Hospital Heidelberg, Medical Faculty Heidelberg, Heidelberg University, 69120 Heidelberg, Germany; German Center for Cardiovascular Research (DZHK), Partner Site Heidelberg/Mannheim, Germany

**Keywords:** Nonsense-mediated mRNA decay, induced pluripotent stem cells, cardiomyocytes

## Abstract

Nonsense-mediated mRNA decay (NMD) serves as a mechanism to suppress the expression of mutant alleles containing premature termination codons, limit the expression of aberrantly spliced transcript isoforms, and control the expression of numerous regular genes. While the principles by which NMD recognizes target transcripts are well understood, much less is known about the range of mRNAs subject to NMD in tissues and specialized cell types. Here we describe the landscape of genes whose expression is controlled by NMD in cardiomyocytes derived from human induced pluripotent stem cells (iPSC-CM), using small read RNA sequencing in combination with a potent inhibitor of SMG1, a kinase essential for NMD. We find that NMD targets are highly conserved between iPSC-CM lines derived from two healthy individuals. Beyond gene level analysis, we identify individual exon and intron RNA sequences that strongly accumulate upon SMG1 inhibition. Using the cardiac NMD targets identified at gene, exon and intron level, we then demonstrate reduced NMD efficiency upon knockdown of two essential NMD factors, UPF1 and UPF2, in iPSC-CM. Our analysis demonstrates that quantifying the transcriptome-wide response to SMG1 inhibition represents a highly sensitive approach to assess global activity of the NMD pathway.

## INTRODUCTION

Nonsense-mediated mRNA decay (NMD) was initially discovered in the context of genetic diseases as a means to suppress the expression of mutated alleles harboring a premature termination codon (PTC) [1, 2]. Since PTCs can lead to the production of potentially harmful truncated polypeptides, degrading the corresponding PTC-containing mRNAs represents a potent strategy to favor expression of the healthy allele and mitigate the severity of such genetic lesions [1, 2]. On the other hand, NMD can also aggravate the phenotypic manifestation of mutations, e.g. when a gene is subject to haploinsufficiency and the truncated polypeptide produced from the PTC-containing allele has the potential to remain functionally active. In addition to its role in suppressing the expression of alleles with inherited or newly acquired PTC mutations, NMD has a major role in silencing transcript isoforms that arise from alternative, spurious or unwanted splicing events that lead to the appearance of a PTC, e.g. though the inclusion of alternative exons or the retention of introns [3, 4]. Here, NMD appears as a quality control mechanism to purify gene loci from transcript isoforms that result from the imperfection of the transcription and splicing machineries. Given that these isoforms quantitatively represent the major group of NMD substrates [3, 4], one may speculate that restricting aberrant transcription and splicing products is the ancestral function of the NMD pathway.

Beyond its role in transcript quality control, NMD was found to be important for regulating the decay rates of regular, protein-coding mRNAs, e.g. during development of the nervous system or in the context of stress responses [1, 2]. Here, NMD appears as a gene regulatory pathway that can shape the dynamics of gene expression and couple mRNA half-lives to the efficiency of protein synthesis. Mechanistically, NMD is thought to be triggered by inefficient or aberrant translation termination events [5]. While regular stop codons are typically located in the last exon of a transcript, aberrant termination occurs, e.g., when a termination codon is located at least 50-55 nt upstream of the most 3’ terminal exon-exon junction, hence referred to as a PTC. During such a translation termination event, the eukaryotic release factors (eRF) 1 and 3 encounter a downstream exon-exon junction complex (EJC), which promotes the assembly of UPF1, UPF2 and UPF3 on the mRNA near the terminating ribosome. This sets off a cascade of events culminating in NMD, including recruitment of the SMG1 kinase, phosphorylation of UPF1 by SMG1, binding of the endoribonuclease SMG6 and the adaptor proteins SMG5/7 to phospho-UPF1, and ultimately endonucleolytic cleave as well as exonculeolytic degradation of the mRNA through recruitment of deadenylating and decapping enzymes [1, 2]. Interestingly, this NMD-activating complex is also assembled at stop codons located in the last exon, e.g. when mRNAs contain upstream open reading frames in the 5’UTR, or when the distance between the stop codon and the poly(A) tail (i.e. the 3’UTR) is exceptionally long. These more common features present in many transcripts explain why several normal protein-coding mRNAs are also subject to the NMD pathway, although they cannot explain the full extent and selectivity of NMD substrate recognition [3, 4].

While substrate recognition and the mechanistic principles of NMD have been intensely studied in recent years, there is still limited understanding of the role of the NMD pathway in specific organs, tissues and cell types. This also holds for the cardiac system, where PTC mutations are known to underlie a substantial proportion of hereditary cardiomyopathies [6], yet physiological targets of NMD are largely unknown. This might be of particular interest in cardiomyocytes, since PTC mutations in the *MYBPC3* gene are not only the most prevalent cause of familial hypertrophic cardiomyopathy [7], but were also proposed to cause chronic activation of NMD [8]. In the study presented here, we made use of the SMG1 kinase inhibitor 11e [9] to characterize the landscape of NMD targets in human induced pluripotent stem cell - derived cardiomyocytes (iPSC-CM). Using the identified sets of cardiac genes, exons and introns that are susceptible to NMD, we further employed RNA expression differences upon SMG1 inhibition as a sensitive and robust measure of the overall NMD pathway activity.

## RESULTS

### Active NMD in human iPSC-derived cardiomyocytes

To assess whether NMD is active in human iPSC-CM, we first made use of a T-cell receptor β(*TCR*β) minigene containing a PTC in the VDJ-exon (Fig. 1A), which had previously been used to study the NMD pathway [10, 11]. Both a wild-type (WT) and a PTC-containing version of the *TCR*β minigene were placed under control of the human Troponin T2 (TNNT2) promoter that is highly active in cardiomyocytes. Upon cloning into an adeno-associated virus (AAV) vector, the minigenes were transduced as AAV6 particles into iPSC-CM line 78 derived from a healthy donor. Expression of *TCR*β-PTC mRNA was reduced more than 10-fold compared to *TCR*β-WT mRNA (Fig. 1B). Moreover, inhibition of SMG1 using the inhibitor 11e [9], henceforth referred to as SMG1-inhibitor (SMG1i), led to a 6-fold increase in *TCR*β-PTC mRNA expression, but only a marginal increase in *TCR*β-WT mRNA expression (Fig. 1C). The results of these reporter gene assays were indicative of active NMD in iPSC-CM.

**Fig. 1.**
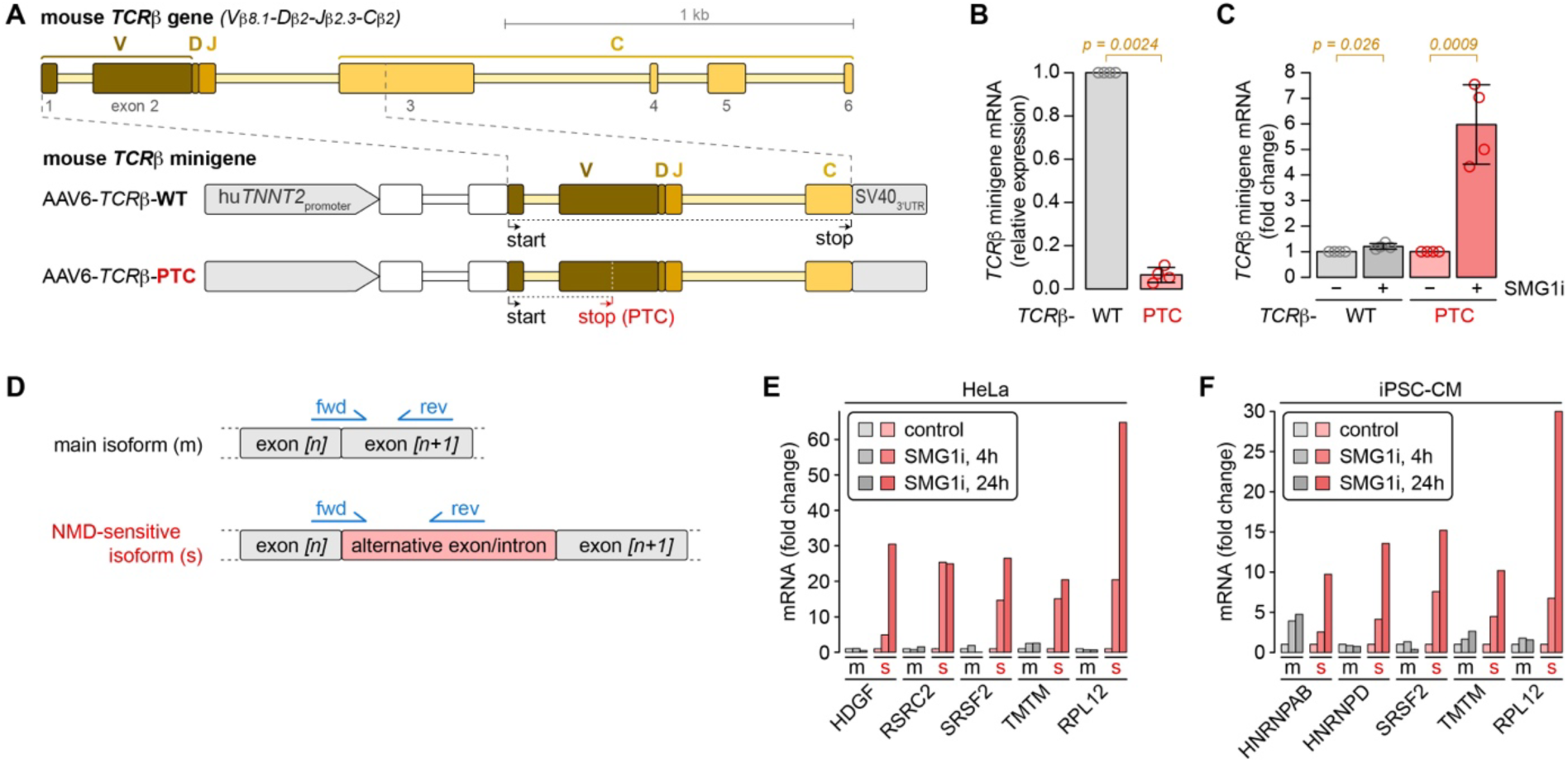
Accumulation of NMD-sensitive transcript isoforms upon SMG1 inhibition in iPSC-CM. **A** Schematic representation of the mouse T-cell receptor β-chain (*TCR*β) gene as well as a *TCR*β minigene, which is flanked by the human troponin T2 (*TNNT2*) promoter and the SV40 3’UTR, and expressed from an AAV vector. Translation of the *TCR*β-WT minigene terminates at a stop codon within the constant region (C), whereas the *TCR*β-PTC minigene terminates at a premature termination codon within the variable region (V). **B** Human iPSC-CM (line 78) were transduced with AAV6-*TCR*β-WT or -PTC particles. After 10 days, cells were treated with DMSO (solvent control) for 24 hours, and total RNA was isolated to determine *TCR*β minigene expression levels by RT-qPCR; values were normalized to the expression level in AAV6-*TCR*β-WT transduced cells (mean ± SD, n = 4 biological repeats, p determined by ratio paired t-test). **C** iPSC-CM (line 78) were transduced as in panel B, and treated with either DMSO (solvent control) or SMG1i (11e, 2.0 μM) for 24 hours. *TCR*β-WT and -PTC minigene expression levels were normalized to the respective DMSO controls (mean ± SD, n = 4 biological repeats, p determined by ratio paired t-test). **D** Schematic illustration of primers (blue) designed to discriminate the main transcript isoform (m) of a gene from a NMD-sensitive isoform (s) generated by inclusion of an alternative exon or a retained intron. **E** HeLa cells were treated for 4 or 24 hours with SMG1i (11e, 1.0 μM), or with DMSO for 24 hours as negative control, followed by isolation of total RNA. RT-qPCR was used to determine the expression levels of m and s transcript isoforms from five genes as indicated. **F** The expression levels of m and s transcript isoforms from five genes were determined in iPSC-CM (line 113) similar to the analysis in panel E.

The same approach was then extended to endogenous genes, whereby we compared the main protein-coding transcript isoform of a gene to a minor transcript isoform of the same gene that is suppressed by NMD, as identified in previous studies [3, 4, 12, 13]. These NMD-sensitive isoforms typically arise though retention of an intron or inclusion of an alternative exon, whereby a PTC is frequently introduced in the transcript. PCR primers were designed to distinguish the main transcript isoform (m, see Fig. 1D) from the NMD-sensitive isoform (s) of 5 genes expressed in the human cervix carcinoma cell line HeLa, and treatment with SMG1i confirmed upregulation of the s-, but not the m-isoforms, for each of the genes tested (Fig. 1E). By the same approach, we then examined 5 genes expressed in cardiomyocytes, using iPSC-CM line 113 derived from a different healthy donor. Again, we observed strong upregulation of the s-isoforms upon SMG1i treatment, whereas the m-isoforms were not, or only mildly, upregulated by SMG1i (Fig. 1F). These experiments confirmed that NMD is active in iPSC-CM, targeting minor transcript isoforms.

### Transcriptome-wide identification of NMD target genes in iPSC-CM

Using iPSC-CM line 113, we then explored the kinetics by which NMD-sensitive (s) transcript isoforms accumulate upon SMG1i inhibition. For four out of the six genes tested, the plateau of s-isoform accumulation was reached after 24 hours of SMG1 inhibition (Fig. 2A). We then tested different concentrations of SMG1i, ranging from 0.5 µM to 10 µM. These dose response curves revealed that for five out of the six genes tested, s-isoform accumulation reached its plateau at a concentration of 2.5 µM SMG1i (Fig. 2B).

**Fig. 2.**
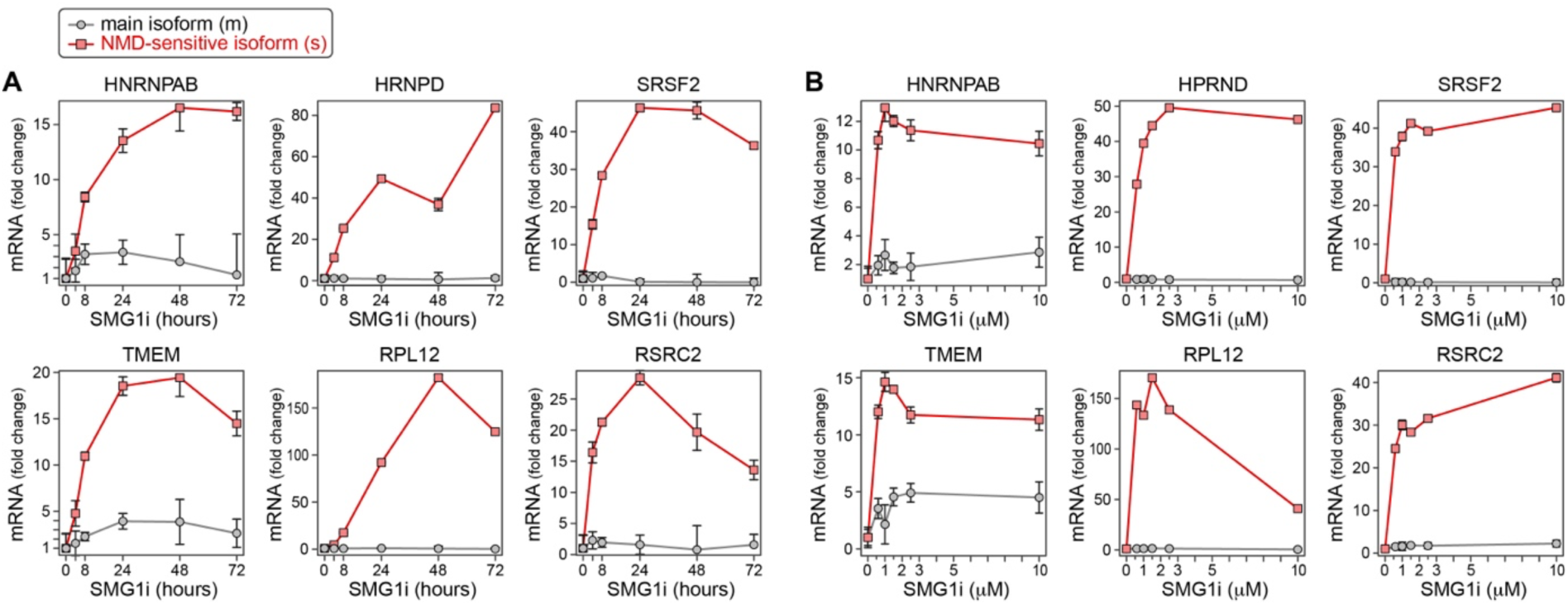
Kinetics and dose response of SMG1 inhibition. **A** Human iPSC-CM (line 113) were treated with SMG1i at a concentration of 2.5 µM for 4, 8, 24, 48, and 72 hours. DMSO treatment for 72 hours served as negative control (shown as 0 hour SMG1i time point). Expression of the main (m) and a NMD-sensitive (s) transcript isoform of *HNRNPAB*, *HNRNPD*, *SRSF2*, *TMEM*, *RPL12*, and *RSRC2* was determined by RT-qPCR. Expression levels were normalized to the expression level of *GAPDH* mRNA, and represented as fold change over the expression level in the DMSO control. Shown are mean values ± SEM, based on n = 3 biological replicates. **B** Human iPSC-CM (line 113) were treated for 24 hours with SMG1i at concentrations of 0.5, 1.0, 1.5, 2.5, and 10 µM. DMSO treatment for 24 hours served as negative control. Expression of the main (m) and a NMD-sensitive (s) transcript isoform of *HNRNPAB*, *HNRNPD*, *SRSF2*, *TMEM*, *RPL12*, and *RSRC2* was determined as in panel A. Shown are mean values ± SEM, based on n = 3 biological replicates.

In the following, we chose iPSC-CM (line 78) to explore the effect of SMG1i on a transcriptome-wide scale. The cells were treated with SMG1i at a concentration of 2.5 µM for 4, 8 and 24 hours, which allowed us to monitor the accumulation of NMD-sensitive transcripts. Following total RNA extraction, small read RNA-Seq was performed on the Illumina NextSeq 2000 platform, using random primers for the reverse transcription step. For the initial analysis, we chose an approach that is agnostic to transcript isoforms by summarizing all reads at the gene level (gene mRNA). This analysis revealed progressive accumulation of approximately 1200 gene mRNAs by 24 hours of SMG1i treatment (red, Fig. 3A-D), whereas approximately 270 gene mRNAs were downregulated by this time (blue).

**Fig. 3.**
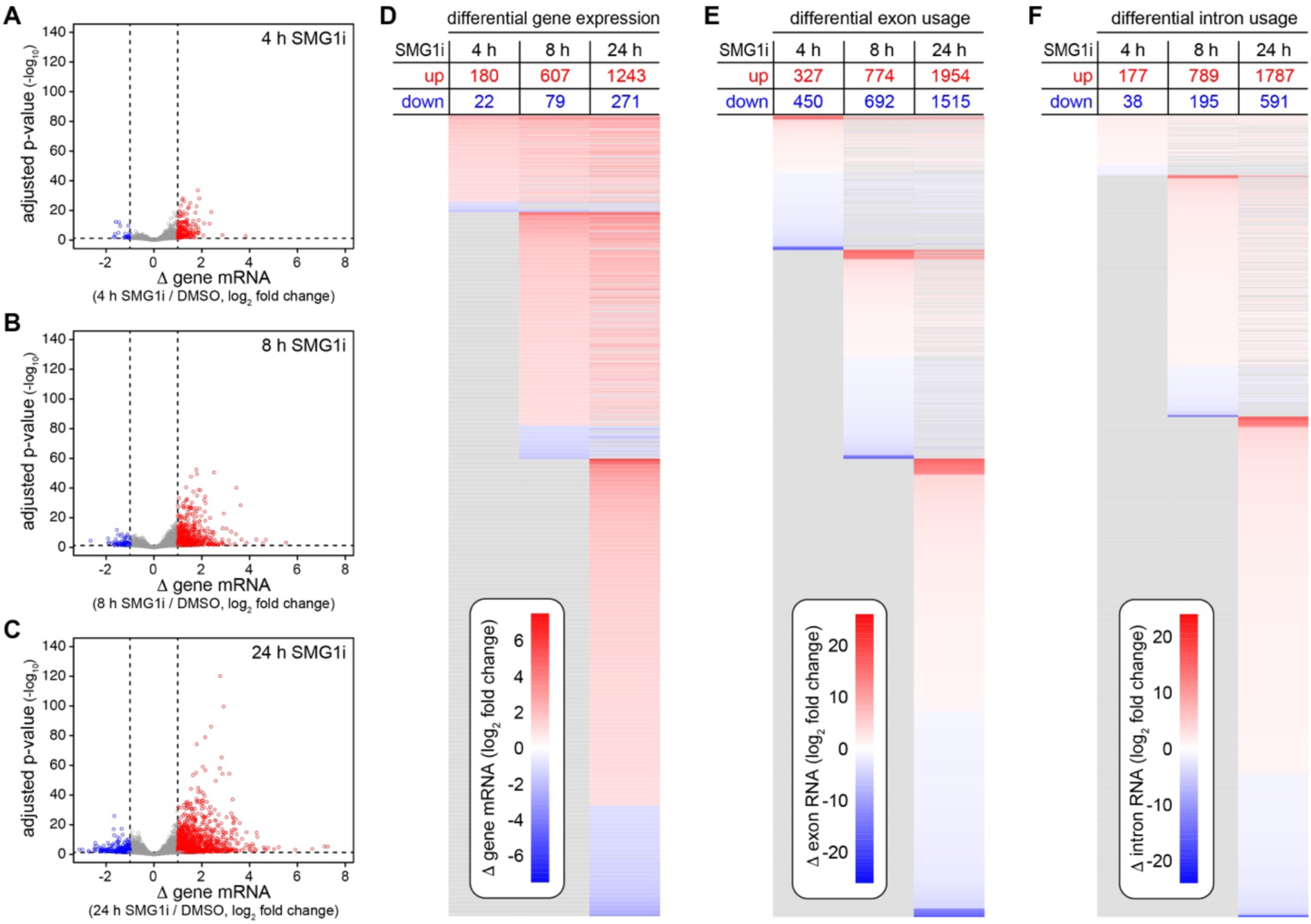
Identification of NMD target RNAs in human iPSC-CM by RNA-Seq. iPSC-CM (line 78) were treated with SMG1i (2.5 µM) for 4, 8, and 24 hours; DMSO treatment for 24 hours served as negative control. Total RNA was depleted of rRNA and subjected to small read RNA-Seq analysis; the experiment was conducted in n = 3 biological repeats. **A** For gene level analysis, all reads mapping to a given gene were summarized and analyzed by DESeq2. The volcano plot depicts the change in gene mRNA expression upon 4 hours SMG1i over the DMSO control, in relation to the p-value adjusted for multiple testing. Genes mRNAs with an adjusted p-value < 0.05 and a change in expression > 2-fold (log_2_ fold change > 1) were considered significantly upregulated NMD targets and depicted in red. Genes mRNAs with an adjusted p-value < 0.05 and a change in expression < 0.5-fold (log_2_ fold change < -1) were considered significantly downregulated and depicted in blue. **B** The same analysis as in panel A was conducted for changes in gene mRNA expression upon 8 hours SMG1i. **C** The same analysis as in panel A was conducted for changes in gene mRNA expression upon 24 hours SMG1i. **D** Heatmap of differentially expressed genes (gene level analysis) upon SMG1i for 4, 8 and 24 hours; the table shows the number of the significantly upregulated and downregulated gene mRNAs. **E** For exon level analysis, the change in usage of individual exons was calculated using DEXSeq. The heatmap depicts differentially used exons after 4, 8 and 24 hours of SMG1i. Exon RNAs with an adjusted p-value < 0.05 and a change in usage > 2-fold (log^2^ fold change > 1) were considered significantly upregulated NMD targets and depicted in red. Exon RNAs with an adjusted p-value < 0.05 and a change in usage < 0.5-fold (log_2_ fold change < -1) were considered significantly downregulated and depicted in blue. **F** For intron level analysis, the same procedure as in panel E was applied to intron RNAs.

### Changes in intron and exon usage reflects NMD-sensitive transcript isoforms

As an alternative approach, we assessed differential usage of individual exons and introns upon SMG1i, which is better suited to reflect the accumulation of NMD-sensitive transcript isoforms. After 24 hours of SMG1i, approximately 1900 exon RNAs and approximately 1700 intron RNAs were found to be upregulated (Fig. 3E and F). Notably, the fold change for some of these exonic and intronic RNA sequences was much higher (> 1000-fold) than the fold change of entire gene mRNAs (reaching 100-fold maximally). This indicates that SMG1i causes pronounced accumulation of numerous aberrant transcript isoforms that are normally under strong suppression of the NMD pathway.

### Validation of NMD targets in a second iPSC-CM line

To validate the NMD targets identified in human iPSC-CM line 78, we made use of iPSC-CM line 113 from another healthy donor, and repeated a similar transcriptome-wide analysis upon treatment with the SMG1i for 24 hours. In this case, we chose a lower concentration of SMG1i, 1.0 µM instead of 2.5 µM, to minimize off-target effects of the kinase inhibitor. The results showed a strong correlation between the global changes in gene expression upon SMG1i in the two iPSC-CM lines, with a Pearson’s correlation coefficient *r* of 0.71 (Fig. 4A). The slope of the regression line (green in Fig. 4A) was below 1, indicating that the effect of SMG1i in line 113 was weaker than in line 78, in line with the lower concentration of SMG1i used in line 113.

**Fig. 4.**
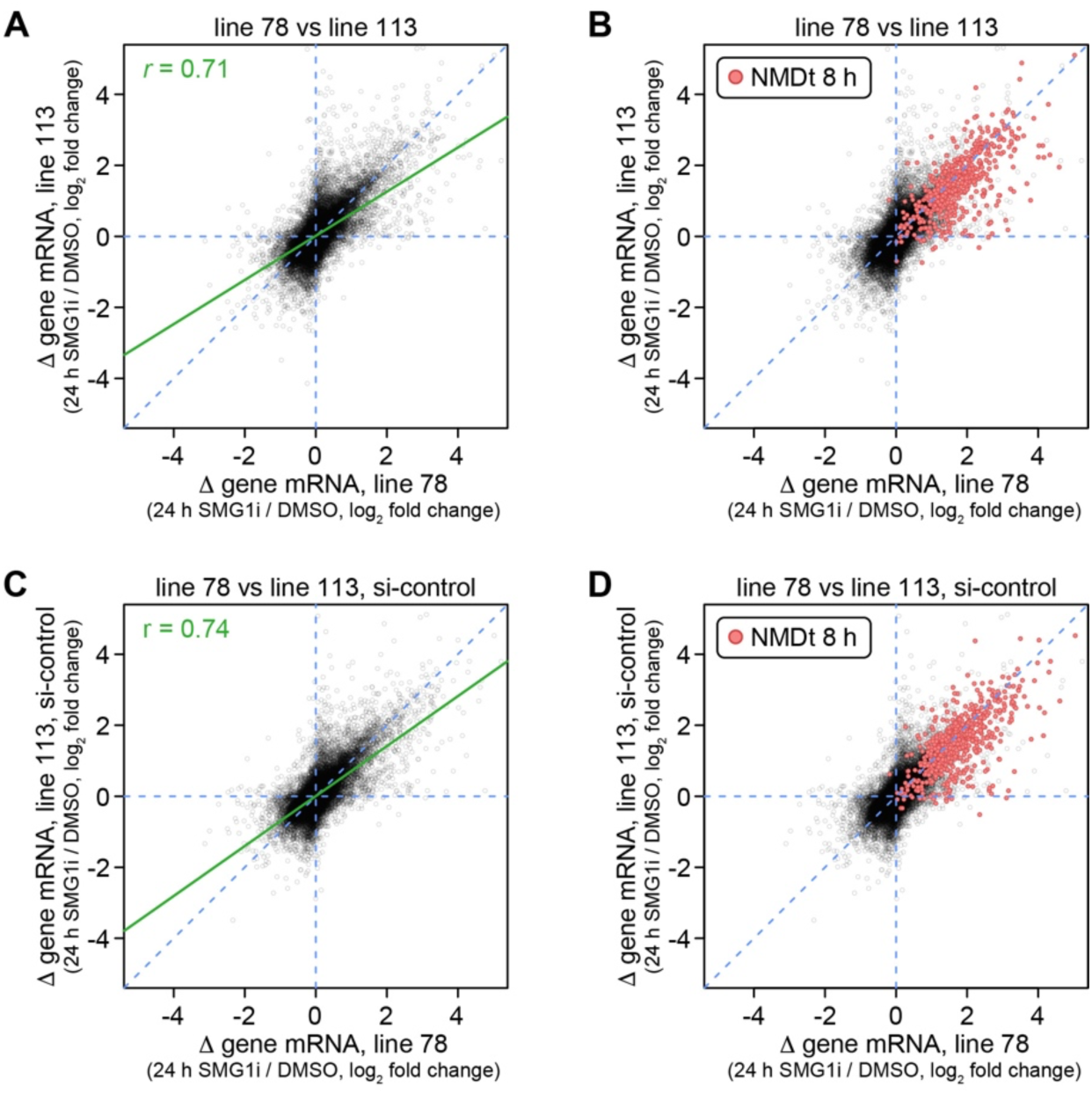
Comparison of NMD inhibition in human iPSC-CM line 78 and line 113. **A** Treatment of iPSC-CM (line 78) with SMG1i (2.5 µM) for 24 hours (data from Fig. 3C) was compared to treatment of untransfected iPSC-CM (line 113) with SMG1i (1.0 µM) for 24 hours; RNA-Seq analysis was conducted as in Fig. 3. The scatter plot shows changes in gene mRNA expression; regression line and Pearson’s correlation coefficient *r* are depicted in green. **B** NMD targets significantly upregulated upon SMG1i treatment for 8 hours in line 78 (identified in Fig. 3B) are shown in red on the scatter plot from panel A. **C** The effect of SMG1i in line 78 (2.5 µM, 24 hours, data from Fig. 3C) was compared to SMG1i treatment (1.0 µM, 24 hours) of iPSC-CM (line 113) transfected with non-targeting control siRNA. The scatter plot shows changes in gene mRNA expression; regression line and Pearson’s correlation coefficient *r* are depicted in green. **D** NMD targets significantly upregulated upon SMG1i treatment for 8 hours in line 78 (identified in Fig. 3B) are shown in red on the scatter plot from panel C.

By visualizing NMD targets as defined by the group of mRNAs significantly upregulated upon 8 hours of SMG1i in line 78 (Fig. 3B), a strong overlap of NMD target mRNAs could be observed between the two iPSC-CM lines (Fig. 4B). This illustrates the conserved nature of the NMD targeting mechanism in iPSC-CM derived from different human donors. In addition, we compared the response of iPSC-CM line 78 cells to 24 hours SMG1i with that of iPSC-CM line 113 cells that had been transfected with a non-targeting control siRNA for 48 hours. Again, the comparison revealed a strong correlation between the global changes in gene expression upon SMG1i (*r* = 0.74, Fig. 4C), and a prominent overlap of NMD target mRNAs (Fig. 4D). Hence, siRNA transfection *per se* does not appear to cause major changes in NMD.

### Knockdown of UPF1 and UPF2 in iPSC-CM

To further interrogate NMD targets in iPSC-CM, we then interfered with NMD activity by knockdown (kd) of two essential NMD factors, UPF1 and UPF2. To this end, iPSC-CM (line 113) were transfected with siRNAs targeting UPF1 or UPF2, or a non-targeting control siRNA. After 48 hours, cells were treated with either SMG1i or DMSO (as solvent control) for 24 hours, followed by small read RNA-Seq analysis. Comparison of normalized read counts for UPF1 and UPF2 mRNA confirmed kd of both mRNAs, whereby UPF1 mRNA kd appeared slightly more efficient (Fig. 5A). Western blot analysis confirmed the successful kd of UPF1 and UPF2 proteins (Fig. 5B).

**Fig. 5.**
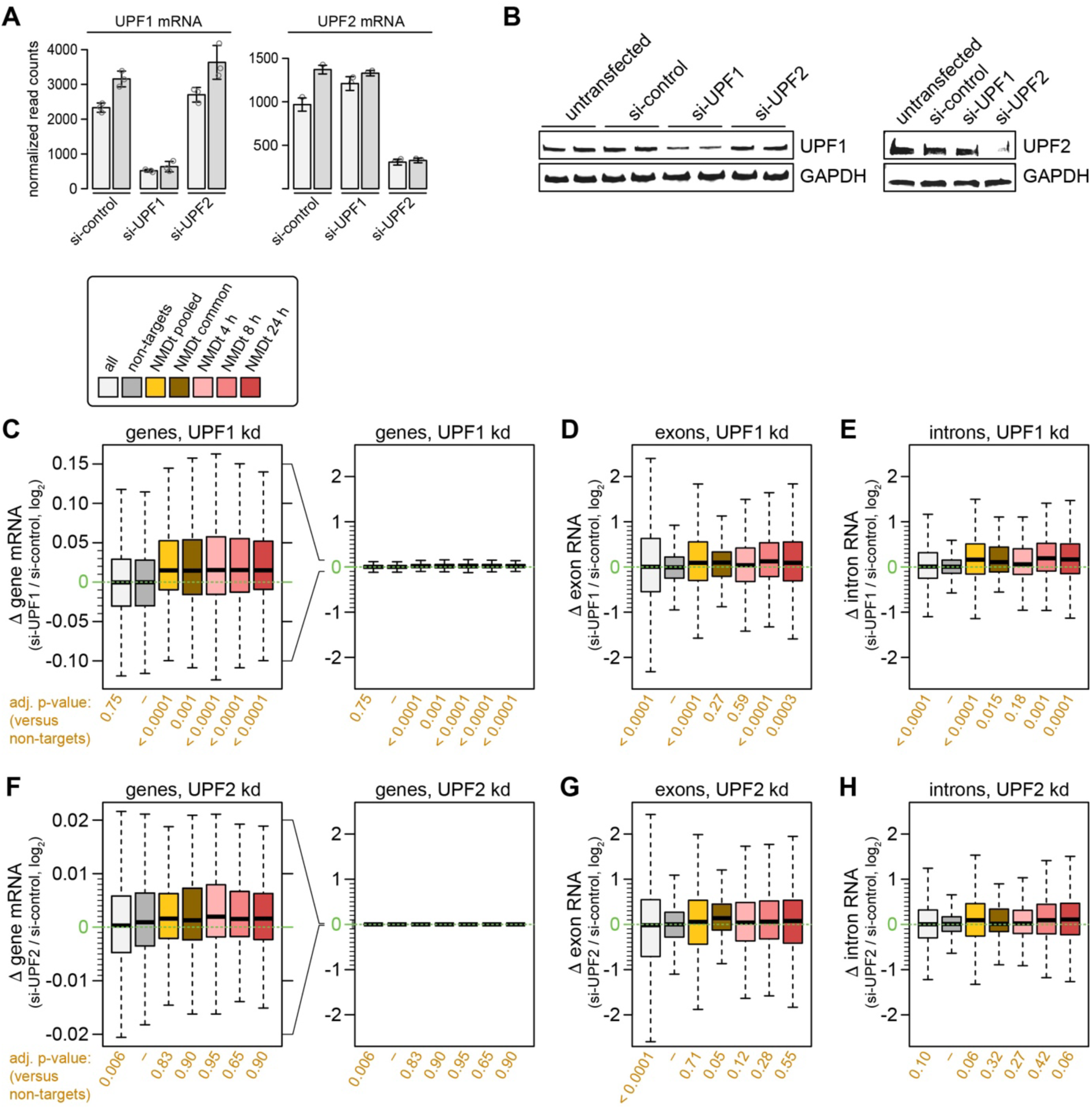
Assessment of NMD activity upon UPF1/2 kd by NMD target RNA expression analysis. **A** UPF1 and UPF2 expression was suppressed by siRNA-mediated knockdown (kd) in iPSC-CM (line 113); a non-targeting siRNA was used as negative control. Subsequently, cells were treated with DMSO for 24 hours. Total RNA was depleted of rRNA and subjected to small read RNA-Seq analysis; the experiment was conducted in n = 3 biological repeats. Normalized read counts for UPF1 and UPF2 mRNA are shown in the barplot (mean value ± SD, n = 3). **B** Western blot analysis of UPF1 and UPF2 was carried out after kd of UPF1 and UPF2 in iPSC-CM (line 113); GAPDH serves as loading control. **C** Changes in gene mRNA expression (gene level analysis) by UPF1 kd are depicted in the boxplot for the following groups defined by their response to SMG1i in iPSC-CM (line 78) from Fig. 3: all genes expressed; non-NMD targets (non-targets) defined by an adjusted p-value > 0.05 and |(log_2_ fold change)| < 0.05 in each of the 4, 8 and 24 hour SMG1i time points; pooled NMD targets (NMDt pooled) encompassing all RNAs significantly upregulated either at the 4, 8 or 24 hour SMG1i time point; common NMD targets (NMDt) restricted to those that are significantly upregulated at all three SMG1i time points; NMDt significantly upregulated at the 4 hour; at the 8 hour; and at the 24 hour SMG1i time point. The boxplot on the right side depicts the same changes in gene mRNA expression by UPF1 kd at the scale of panels D and E. The same categories were applied to depict **D** changes in exon RNA usage by UPF1 kd; **E** changes in intron RNA usage by UPF1 kd; **F** changes in gene mRNA expression by UPF2 kd; **G** changes in exon RNA usage by UPF2 kd; and **H** changes in intron RNA usage by UPF2 kd. Each group was compared to the non-NMD target (non-target) group; corresponding p-values based on a t-test and adjusted for multiple testing are shown below the boxplots.

As a fist approach to assess NMD activity in the UPF1/2 kd iPSC-CM (line 113), we compared RNA levels between the DMSO-treated cells. Prior to the analysis, gene mRNAs, exon RNAs and intron RNAs were grouped according to the response to SMG1i observed in iPSC-CM (line 78, Fig. 3). At the gene mRNA level, no difference could be observed between control kd and UPF1 kd cells when mRNAs from all genes expressed were analyzed as a group (Fig. 5C, light grey), and the same was true for a group of strictly defined non-NMD targets (non-targets, dark grey). In contrast, pooled NMD targets (NMDt pooled, encompassing all RNAs significantly upregulated either at the 4, 8 or 24 hour SMG1i time point in line 78 iPSC-CM) were expressed at higher levels in UPF1 kd cells (Fig. 5C, yellow). The same was observed for the group of common NMD targets (significantly upregulated at all three SMG1i time points, brown) and for the groups of NMD targets significantly upregulated at the 4 hour (pink), 8 hours (light red) and 24 hour SMG1i time point (dark red). While the elevated expression of these groups in UPF1 kd cells was statistically significant in all cases (p-values of 0.001 and below), the magnitude of elevated expression was small (median log_2_ fold change approximately 0.015).

When the same analysis was conducted with exon (Fig. 5D) and intron usage (Fig. 5E), we also observed elevated expression of NMD target exons and introns in the UPF1 kd cells. Here, the magnitude of elevated intron and exon expression was about 10 times larger (median log_2_ fold change in the range of 0.1–0.2), yet statistical significance was only reached with the groups of pooled, 8 hour and 24 hour NMD targets. Interestingly, exon and intron usage in general showed much greater changes, visible as larger boxes and whiskers in the boxplots (compare right panel of Fig. 5C to Fig. 5D and E, with identical scales). These strong changes in exon and intron usage reflect pronounced variation in the expression of transcript isoforms.

Knockdown of UPF2 gave a similar pattern, yet the magnitude of elevated gene mRNA, exon RNA and intron RNA expression was smaller for all NMD target groups, and did not reach statistical significance except for one group, common NMD target exons (Fig. 5F-H). This result indicated that kd of UPF2 had a smaller effect on NMD activity, possibly due to a lower kd efficiency compared to UPF1.

### Response to SMG1 inhibition allows sensitive assessment of global NMD activity

As a second approach to assess NMD activity, we compared the effect of SMG1i between the control kd and UPF1 kd in iPSC-CM (line 113). At the gene level analysis, upregulation of NMD target gene mRNA expression upon SMG1i was less pronounced in the in the UPF1 kd cells than in the control kd cells, visible by a slope of the regression line below 1 (red line in Fig. 6A). Reduced upregulation of mRNA expression was observed for all NMD target gene groups in UPF1 kd cells, with high statistical significance (Fig. 6B). When the same analysis was conducted for differences in exon and intron usage, a significantly reduced upregulation in UPF1 kd cells was observed for three of the five NMD target exon groups (Fig. 6C), and four of the five NMD target intron groups (Fig. 6D). Interestingly, statistical significance was generally higher with the intron analysis, indicating that intron usage offers a more robust read-out for NMD activity than exon usage.

**Fig. 6.**
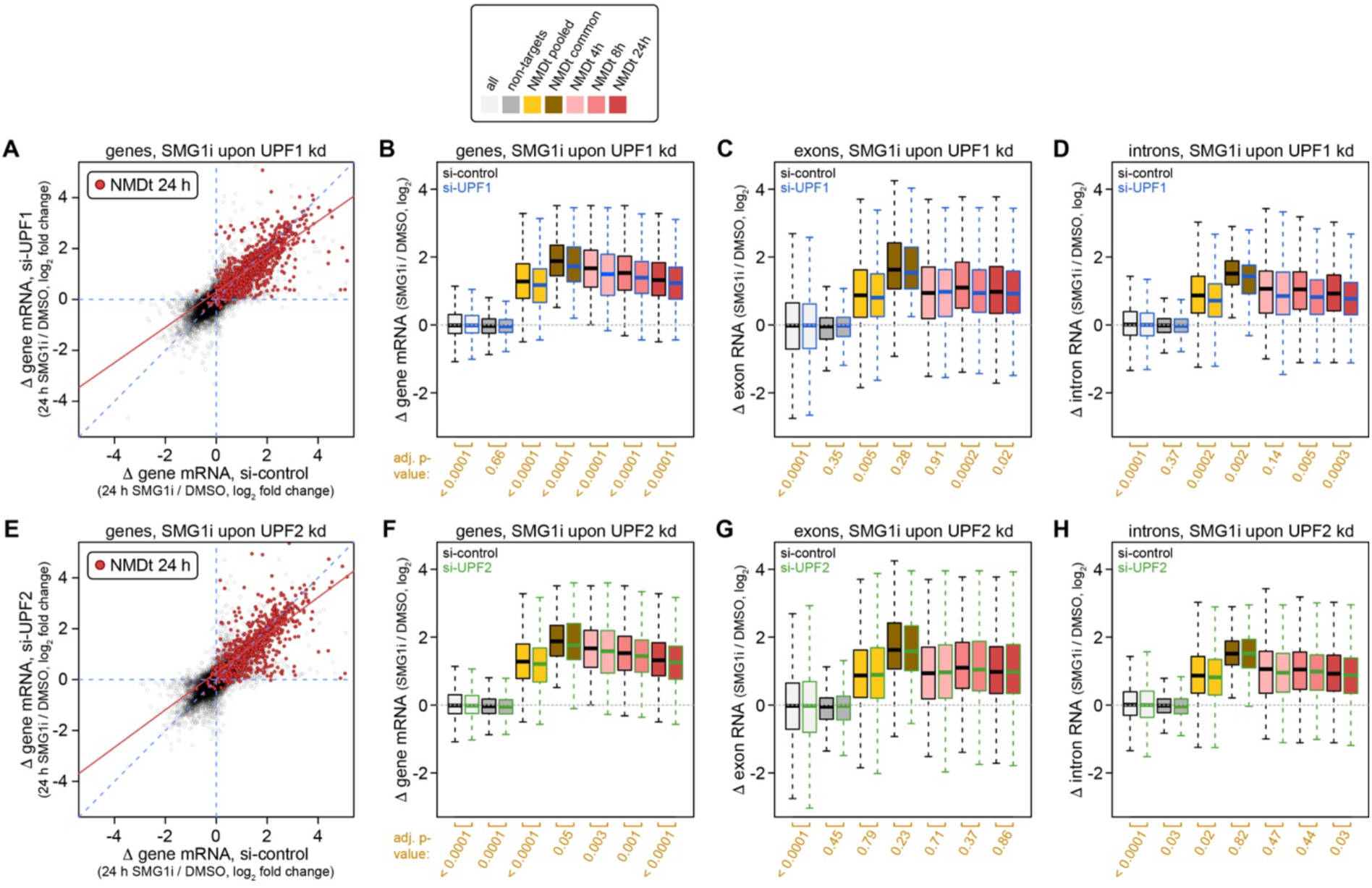
Assessment of NMD activity upon UPF1/2 kd by comparison of the SMG1i effect. Human iPSC-CM (line 113) were transfected with siRNAs targeting UPF1 or UPF2, or with non-targeting control siRNA, as in Fig. 5. After 48 hours, cells were treated with SMG1i (1.0 µM) or DMSO as solvent control for 24 hours. RNA-Seq analysis was conducted as in Fig. 3; the experiment was carried out in n = 3 biological repeats. **A** The scatter plot shows changes in gene mRNA expression upon SMG1i, comparing UPF1 kd with control kd; NMD targets (NMDt) significantly upregulated upon SMG1i treatment for 24 hours in line 78 (identified in Fig. 3C) are shown in red, together with the regression line for these NMD targets. **B** Changes in gene mRNA expression (gene level analysis) by SMG1i are depicted in the boxplot for groups defined by their response to SMG1i in iPSC-CM line 78 (data from Fig. 3), as introduced in Fig. 5C. The SMG1i effect is compared between control kd and UPF1 kd cells. **C** Changes in exon RNA usage upon SMG1i, comparing the SMG1i effect between control kd and UPF1 kd cells. **D** Changes in intron RNA usage upon SMG1i, comparing the SMG1i effect between control kd and UPF1 kd cells. **E** The scatter plot shows changes in gene mRNA expression upon SMG1i, comparing UPF2 kd with control kd, as in panel A. **F** Changes in gene mRNA expression upon SMG1i, comparing the SMG1i effect between control kd and UPF2 kd cells. **G** Changes in exon RNA usage upon SMG1i, comparing the SMG1i effect between control kd and UPF2 kd cells. **H** Changes in intron RNA usage upon SMG1i, comparing the SMG1i effect between control kd and UPF2 kd cells. For each comparison, p-values based on a paired t-test and adjusted for multiple testing are shown below the boxplots.

When the gene level analysis was applied to the UPF2 kd cells, we also observed reduced upregulation of NMD target gene mRNA expression upon SMG1i, again visible by a slope of the regression line below 1 (Fig. 6E). The tilt in the regression line was less than that observed with the UPF1 kd cells, confirming that the UPF2 kd condition caused a weaker inhibition of NMD activity than the UPF1 kd. Again, a significantly reduced upregulation of gene mRNA expression was observed for all NMD target gene groups in UPF2 kd cells (Fig. 6F), though with lower statistical significance than in UPF1 kd cells. Exon and intron usage analysis showed barely any difference in the SMG1i response between control kd and UPF2 kd cells (Fig. 6G and H), with the exception of significantly reduced upregulation of pooled NMD target introns (Fig. 6H, yellow) and 24 hour NMD target introns (dark red). Thus, the UPF2 kd condition revealed that the analysis of SMG1 inhibition effects is more sensitive for measuring changes in NMD activity than the simple analysis of NMD target RNA levels.

## Discussion

Given its broad role in mitigating the consequences genetic nonsense mutations, suppressing aberrantly spliced transcript isoforms and controlling the expression of specific protein-coding mRNAs, NMD is critical for maintaining cellular homeostasis through RNA quality control and shaping gene expression programs. It is therefore not surprising that components of the NMD machinery were found to be essential for organismal development [14], as exemplified by loss of UPF1 and 2 causing early embryonic lethality in both zebrafish [15] and mice [16, 17].

Interestingly, NMD activity itself was found to vary between different human tissues [18], and there is evidence that activity of the NMD machinery is altered in specific diseases including distinct types of cancer [19]. Reduced NMD activity, e.g., was observed in pancreatic adenosquamous carcinoma due to frequent UPF1 mutations [20], whereas enhanced NMD activity was reported in colorectal cancers with microsatellite instability [21]. Chronic activation of NMD was also proposed to occur in a specific subtype of genetic hypertrophic cardiomyopathy that is driven by PTC mutations in the *MYBPC3* gene [8]. These findings raise the need for identifying tissue- and cell type-specific targets of NMD, and for developing methods to faithfully assess the overall activity of the NMD machinery in such systems.

Here we chose iPSC-CM derived from two healthy human donors as an experimentally tractable system for identifying cardiac NMD targets and measuring NMD activity. Using a heterologous NMD reporter based on a murine *TCR*β minigene, as well as a small set of endogenous NMD-sensitive transcript isoforms, we show that NMD is active in iPSC-CM (Fig. 1). Time course and dose response experiments allowed us to determine conditions where NMD is fully inactivated by the SMG1 inhibitor 11e, and NMD target accumulation has reached its plateau (24 hours, 1.0 – 2.5 µM SMG1i, Fig. 2). These conditions were then used to determine NMD target RNAs on a transcriptome-wide scale in the iPSC-CM line 78 using a differential gene mRNA expression approach as well as a differential exon and intron usage approach (Fig. 3). NMD targets were also determined in iPSC-CM line 113 from a separate healthy donor, and direct comparison between the two lines revealed an extensive overlap of NMD target genes (Fig. 4), indicating that NMD targeting features are highly conserved between individuals.

Finally, we tested two methods to assess overall NMD activity in iPSC-CM where NMD activity was reduced by UPF1 and UPF2 kd. In the first and more simple approach, the expression levels of NMD target RNAs were directly compared between control kd and UPF1/2 kd cells (Fig. 5). This analysis was able to faithfully detect reduced NMD activity upon UPF1 kd by gene expression analysis with each of the NMD target groups (Fig. 5C). Reduced NMD activity was also observed by exon and intron usage analysis, yet only with those groups containing larger numbers of NMD targets (NMDt pooled, NMDt 8 h, NMDt 24 h; Fig. 5D and E). Notably, intron usage analysis gave larger fold changes and more reliable results than exon usage analysis, which might be due to the fact that introns are typically longer than exons, thus generating more reads from transcript isoforms with retained introns. In contrast to UPF1 kd, the first approach could not faithfully detect reduced NMD activity upon UPF2 kd (Fig. 5F-H).

A similar approach for the assessment of NMD activity was recently applied to a large range of publicly available human RNA datasets, making use of expression differences of endogenous NMD target genes [18]. This study was able to identify genetic alterations in genes encoding NMD factors that correlate with differences in overall NMD activity, assess differences in NMD activity between tissues and individuals, and further extend the methodology to allele-specific expression differences based on the occurrence of NMD-triggering PTCs [18].

In the second approach, we quantified the changes in gene expression upon SMG1i, thereby measuring the difference between the normal state, where RNAs are subject to the actual NMD activity, and a state of full NMD inhibition. Applied to gene mRNA expression, this analysis was able to faithfully detect reduced NMD activity for both UPF1 and UPF2 kd cells (Fig. 6A, B, E and F), indicating that the second approach is more sensitive than the first approach.

The second approach also detected reduced NMD activity in the UPF1 kd cells by both exon and intron usage analysis, though statistically significant differences were again observed only with groups containing larger numbers of NMD targets (NMDt pooled, NMDt 8 h, NMDt 24 h; Fig. 6C and D). In the UPF2 kd cells, exon/intron usage analysis revealed reduced NMD activity only with two intronic NMD target groups (NMDt pooled and NMDt 24 h; Fig. 6H). Thus, the second approach also has a slight advantage when applied to intron and exon usage. Taken together, quantifying the transcriptome-wide response to SMG1 inhibition appears to be a highly sensitive approach for assessment of differences in overall NMD activity. We propose to use such methodology for the quantification of NMD activity in disease models, and expect novel insight to come out of improved assessment of the RNA turnover machinery.

## Materials and Methods

### Cell culture

HeLa cells were cultured in Dulbecco’s modified Eagle’s medium (DMEM, Gibco) supplemented with 10% fetal bovine serum (FBS, Sigma), 2 mM L-glutamine, 100 U/ml penicillin and 0.1 mg/ml streptomycin (all PAN Biotech) in a humidified incubator at 37 °C and 5% CO_2_.

Human iPSCs were cultured in feeder-free and serum-free conditions in DMEM/F12 medium (Corning, 15-090-CM) supplemented with homemade E8 supplement on Matrigel-coated plates (growth factor-reduced, Corning, 354230) in a humidified incubator at 37 °C and 5% CO_2_.

Homemade E8 supplement was prepared by combining: L-ascorbic acid 2-phosphate (64 mg/l, Sigma-Aldrich, A92902-25G), Insulin (19.4 mg/l, Life Technologies, 17504-001), transferrin (10.7 mg/l, Sigma-Aldrich, T8158-100MG), sodium selenite (14 µg/l, Sigma-Aldrich, S5261-10G), FGF2 (100 µg/l, Life Technologies, 100-18B-50UG) and TGFβ1 (2 ng/l, Life Technologies, 100-21-10UG).

Differentiation of human iPSC into CMs was performed using a small-molecule Wnt-activation/inhibition protocol [22]. For the first 7 days, human iPSCs were differentiated using cardiac differentiation medium (CDM3) composed of RPMI1640 (Life Technologies, 21875091) supplemented with L-ascorbic acid 2-phosphate (0.213 g/l, Sigma-Aldrich, A92902-25G) and recombinant human albumin (0.5 g/l, Sigma-Aldrich, A9511-100MG) in sterile water. From day 1-3, the iPSCs were cultured in CDM3 medium containing a gradient concentration of CHIR99021 (4-2 µmol/l, TOCRIS, 4953). From day 3-5, media was changed to CDM3 medium supplemented with IWP2 (5 µmol/l, Med Chem Express, 686770-61-6), followed by CDM3 only from day 5-7. From day 7 onward, cells were cultured in RPMI1640 supplemented with B27 (Life Technologies, 17504044). From day 10-13, hiPSC-CM were maintained in RPMI1640 without glucose, supplemented with CDM3 supplement and lactate. At day 13-15, beating human iPSC-CM were either cryopreserved using Cryo-Brew freezing medium (Miltenyi Biotec, 130-109-558), or passaged using TrypLE select enzyme (Life Technologies, A1217702) followed by resuspension in passaging medium RPMI1640 containing B27, knockout serum replacement (Life Technologies, 10828028) and Rock inhibitor (Sigma-Aldrich, Y0503-1MG), and reseeding on Matrigel-coated plates. All downstream experiments were conducted using human iPSC-CM with >90% purity (evaluated by flow cytometry using Cardiac Troponin T Antibody, Miltenyi Biotec, 130-119-575) between day 45 and 50 after the initiation of differentiation.

### Plasmid cloning

The WT control and PTC-containing mouse *TCR*β minigenes were amplified by PCR from plasmids EV107-433 (WT) and EV107-434 (PTC) kindly provided by Oliver Mühlemann (University of Bern, Switzerland). For this, the following primers were used:

forward: 5’- GATCGATCGGTACCTCGACAGGGCTGGAACAA-3’

reverse: 5’- GATCGATCCTCGAGCCCCGGGCTGCAGGC-3’

The PCR amplicon was then subjected to Sanger sequencing for validation, and ligated into the recombinant AAV backbone vector ST1-pSSV9-hTNTv using *Kpn*I and *Xho*I restriction sites, resulting in plasmids pSSV9-hTNT-*TCR*β-WT and pSSV9-hTNT-*TCR*β-PTC. After cloning, the final plasmids were confirmed by Oxford Nanopore DNA sequencing.

### Production of AAV6 particles

For preparation of recombinant AAV6 particles, we used the procedure as described by Doroudgar et al [23], with slight modifications: HEK293T cells were plated in T-175 flask and maintained in DMEM containing high glucose and pyruvate (Thermo Fischer Scientific, #21969035), supplemented with 10% FBS. Twenty-four hours after plating, cultures were transfected using polyethyleneimine (0.52 mg/ml) in serum-free DMEM with plasmids pSSV9-hTNT-*TCR*β-WT or pSSV9-hTNT-*TCR*β-PTC together with the AAV6 helper plasmid pDP6rs [24]. Twenty-four hours after transfection, medium was changed to DMEM containing 0.5% FBS. Three days later, the cells were collected, centrifuged at 500 g for 10 minutes and resuspended in 10 ml lysis buffer (150 mM NaCl, 50 mM Tris-HCl). The resuspended cells were then subjected to three rounds of freeze-thawing, followed by treatment with Benzonase (1500 U, Sigma Aldrich E1014). Cell debris was collected by centrifugation at 3,400 g for 20 minuntes. The supernatant containing the AAV6 particles was then precipitated using 2.38 M ammonium sulfate, and further purified on an iodixanol gradient comprised of the following four phases: 7.3 ml of 15%, 4.9 ml of 25%, 4 ml of 40%, and 4 ml of 60% iodixanol (Optiprep) overlaid with 10 ml of cell supernatant. The gradients were centrifuged at 50,000 rpm for 2 hours at 4 °C using OptiSeal Polyallomer Tubes (Beckman) in a 50.Ti Rotor of a Beckman ultracentrifuge. Viral particles were collected by inserting a needle 2 mm below the 40–60% interface and collecting four or five fractions (∼4 ml) of this interface and most of the 40% layer. The fractions were analyzed for viral particle content and purity by examining 10 μl of each fraction on a 12% SDS-polyacrylamide gel, followed by staining with Coomassie Blue (Instantblue, Sigma-Aldrich) to visualize the viral capsid proteins VP1, VP2, and VP3. Viral particles were then collected from the fractions of several gradients, and the buffer was exchanged with Ringer’s lactate solution (Braun) using an ultrafiltration device. The final AAV6 particle preparation was then resolved on a 12% SDS-polyacrylamide gel and stained with Instantblue.

### siRNA-mediated knockdown

Knockdown experiments were performed using Lipofectamine RNAiMax (Life Technologies) according to the manufacturer’s instructions. Human iPSC-CMs were transfected with either non-targeting control siRNA (Horizon Discovery, D-001810-01-05), or custom siRNAs targeting either UPF1 (si-hUPF1; 5’-GAUGCAGUUCCGCUCCAUU-dTdT-3’) or UPF2 (si-hUPF2; 5’-GCUCGGAAUUUUUAUGAGA-dTdT-3’) (both from Horizon Discovery Dharmacon at a final concentration of 10 nM for 48 hours prior to subsequent downstream analysis.

### SDS-polyacrylamide gel electrophoresis and western blot analysis

For protein extraction, human iPSC-CMs were lysed in RIPA buffer (Life Technologies, 89900) containing phosphatase and protease inhibitors (Life Technologies, 78442) for 10 minutes. Supernatant was collected after centrifugation at 14,000 g for 20 minutes at 4 °C. Protein concentration was determined by BCA assay (Life Technologies, 23227) according to the manufacturer’s instructions. Samples were mixed with 4× Laemmli Buffer (Bio-Rad, 1610747) supplemented with 10% 2-mercaptoethanol (Sigma-Aldrich, M3148-25ML) and denatured at 95 °C for 5 minutes. 10 µg of protein was loaded onto a 4-15% Mini-PROTEAN TGX precast gels (Bio-Rad, 4561084) and separated by SDS-polyacrylamide gel electrophoresis (80 V for 20 minutes, then 120 V for 50 minutes). Afterwards, proteins were transferred to a nitrocellulose membrane (Bio-Rad, 1610376) using the Trans-Blot Turbo Transfer System (Bio-Rad) at 25 V and 1.3 A for 7 minutes, and stained with Ponceau-S(Sigma-Aldrich, P3504-10G) for 5 minutes to confirm efficient protein transfer.

Membranes were blocked for 1 hour in blocking solution (5% milk in TBS-T, Carl Roth, T145.3) and incubated overnight at 4 °C with the following primary antibodies: mouse monoclonal anti-GAPDH (0411, Santa Cruz Biotechnology, sc-47724, 1:1000), rabbit monoclonal anti-UPF1 (D15G6, Cell signaling, 12040, 1:1000) and rabbit monoclonal anti-UPF2 (D3B10, Cell signaling, 11875, 1:1000). Membranes were washed three times with TBS-T before applying the secondary donkey anti-mouse Alexa Fluor Plus 488 (Life Technologies, A32766, 1:2000) and donkey anti-rabbit Alexa Fluor Plus 647 (Life Technologies, A32795, 1:2000) antibodies at room temperature for 1 hour, protected from light. After washing with TBS-T, fluorescence signals were detected using an Amersham Laser Scanner. Signal intensities were quantified using ImageJ software and the resulting data were plotted using GraphPad Prism 10.1.1.

### RT-qPCR analysis

Cells were collected and lysed in Trizol (Thermo Fisher Scientific), and RNA was extracted with the RNA Clean & Concentrator Kit (Zymo Research) following the manufacturer’s protocol. On-column DNase digestion was performed to remove genomic DNA using DNase I (Zymo Research #E1011). One μg of total RNA was used for cDNA synthesis with the 5× PrimeScript RT Master Mix kit containing both oligo-dT and random primers (Takara, #RR036). RT-qPCR reactions were assembled in 384-well plates using a 5-fold diluted cDNA reaction, 2 μM of each isoform-specific primer, and the TB Green Premix Ex Taq containing ROX reference dye (Takara #RR420W) in a final volume of 10 μl per well. RT-qPCR was performed on a QuantStudio 5 system (Thermo Fisher Scientific). Triplicates were measured for every isoform and the GAPDH reference mRNA. Negative (minus RT) controls, where the reverse transcriptase was heated and inactivated prior to cDNA synthesis, were included in all measurements to verify efficient removal of genomic DNA. Primers were synthesized by Eurofins Genomics, sequences are listed in Table 1.

**Table 1.**
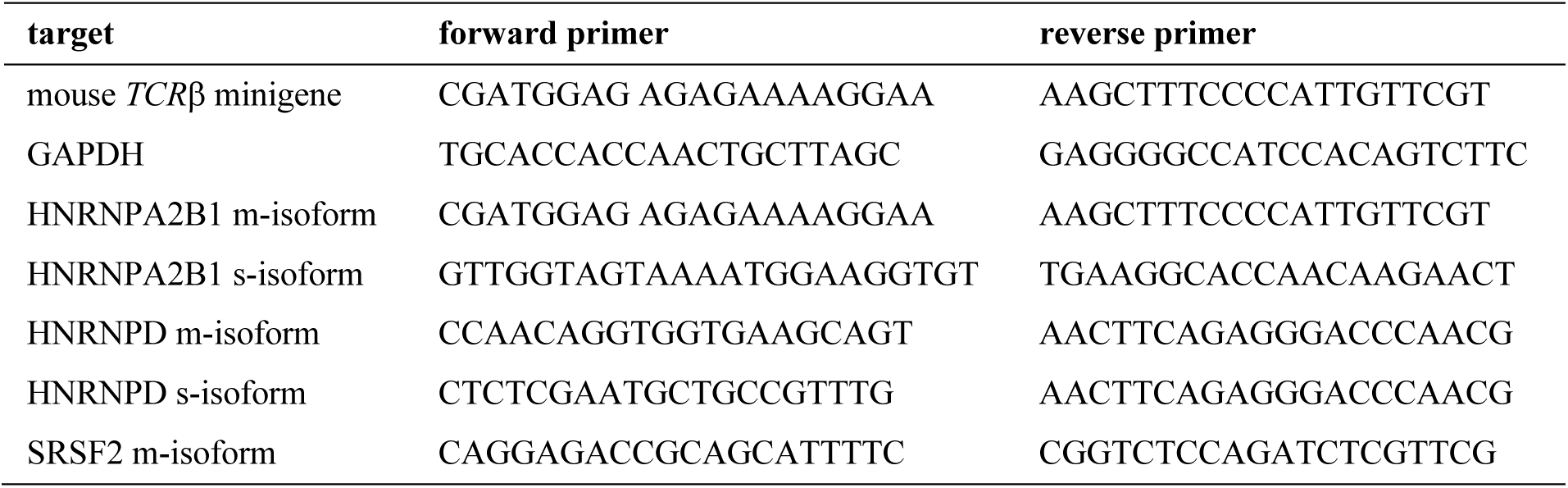
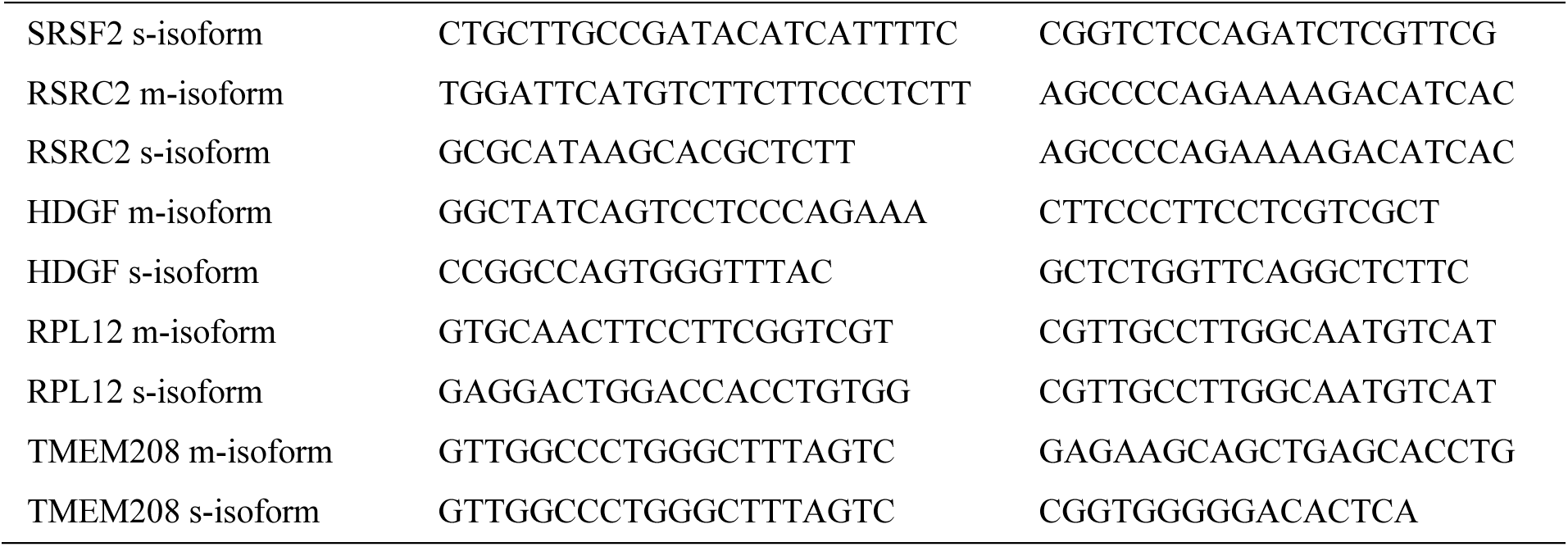
Primer sequences used for RT-qPCR analysis, 5’ to 3’.

### RNA-Seq

RNA-Seq was carried out with two human iPSC-CM lines, line 78 and line 113. Line 78 was analyzed under the following 4 experimental conditions: DMSO-24h, SMG1i-4h, SMG1i-8h, and SMG1i-24h, with three biological replicates each; 12 samples in total. Line 113 was analyzed under the following 8 experimental conditions: untransfected-DMSO-24h, untransfected-SMG1i-24h, si-control-DMSO-24h, si-control-SMG1i-24h, si-UPF1-DMSO-24h, si-UPF1-SMG1i-24h, si-UPF2-DMSO-24h, and si-UPF2-SMG1i-24h, with three biological replicates each; 24 samples in total. Cells were collected and lysed in Trizol (Thermo Fisher Scientific), and RNA was extracted with the RNA Clean & Concentrator Kit (Zymo Research). On-column DNase digestion was performed during RNA purification using DNase I (Zymo Research #E1011). Quality control RT-qPCR for several NMD-sensitive isoforms with including minus RT as negative control was performed prior to library preparation.

One μg total RNA was used for rRNA depletion using the Ribo-Zero Plus rRNA depletion kit (Illumina), and 10 ng of the rRNA-depleted RNA was used for preparing libraries with the NEBNext Ultra II Directional RNA Library Kit (New England Biolabs), which contains random primers for reverse transcription. After quality assessment of the libraries with an Agilent 2100 Bioanalyzer, samples were pooled at equimolar ratio for multiplexing, and sequenced on an Illumina NextSeq 2000 system using the protocol for 100-base single-end reads.

### Bioinformatics analysis

Reads were trimmed using the Trimmomatic tool (version 0.36), thereby removing low-quality and adapter sequences from FASTQ files [25]. Quality of the trimmed FASTQ files was then assessed using FastQC (version 0.11.5), and reads were mapped with STAR (version 2.7.11b) to a primary assembly of the human genome reference (GRCH38) using Homo_sapiens.GRCH38.115.gtf as the annotation file with the default options of STAR [26].

For analysis at the gene expression level, the featureCounts function of the subread package v1.6.395 was used to summarize the read counts for each gene [27]. Differential gene expression analysis was performed with the DESeq2 R-package (1.28.1) [28]. Genes with fewer than 10 read counts across each group were removed. A gene was considered differentially expressed if the p-value adjusted for multiple testing using the Benjamini-Hochberg method was < 0.05 and the absolute fold change in expression was > 2 (log_2_ fold change > 1).

For differential exon and intron usage analysis, a flattened annotation file for exons was generated from the Homo_sapiens.GRCH38.115.gtf file using the GenomeRanges and DEXSeq packages in R [29]. The flattened annotation file for exons served as the basis for the annotation file for introns. Differential exon and intron usage analysis was performed using the DEXSeq library [30, 31]. Analyses were conducted separately for exons and introns. Exons and introns with a p-adj value < 0.05 and an absolute log_2_ fold change > 1 were considered differentially used exons and introns.

### Statistical analysis

Data analysis was performed using R and python. Heat maps were generated using the pheatmap library in R. Student’s t-test was used to compare differences between 2 groups, a paired t-test was used to compare values between two conditions (e.g. si-UPF1 versus si-control). All data in bar graphs are presented as mean ± SD, unless indicated otherwise.

## Acknowledgments

We thank Oliver Mühlemann (University of Bern, Switzerland) for generously sharing NMD reporter gene plasmids as well as for helpful discussions on the project. The human iPSC lines were a kind gift from Joseph C. Wu (Stanford Cardiovascular Biobank, California, USA) to T.S. We are grateful to David Ibberson from the Deep Sequencing Core Facility at Heidelberg University for support with next generation sequencing. This work was funded by the Deutsche Forschungsgemeinschaft (DFG) project numbers 464424253-CRC 1550 to G.S., T.S. and M.V.; 278001972-TRR 186 to G.S.; 439669440-TRR 319 to G.S., 445549683-RTG 2727 to G.S and J.S.; and 462241601 to T.S.

## Author contributions

M.N. and A.A. performed the majority of experiments; R.K., D.L. and V.K.S. supported experimental procedures; J.S. assisted in statistical analyses; M.V., T.S. and G.S. conceived the study; M.N., A.A., T.S. and G.S. interpreted the results and wrote the manuscript.

## References

1. Lykke-Andersen S, Jensen TH: Nonsense-mediated mRNA decay: an intricate machinery that shapes transcriptomes. Nat Rev Mol Cell Biol 2015, 16:665–677.

2. Kurosaki T, Popp MW, Maquat LE: Quality and quantity control of gene expression by nonsense-mediated mRNA decay. Nat Rev Mol Cell Biol 2019, 20:406–420.

3. Karousis ED, Gypas F, Zavolan M, Muhlemann O: Nanopore sequencing reveals endogenous NMD-targeted isoforms in human cells. Genome Biol 2021, 22:223.

4. Boehm V, Wallmeroth D, Wulf PO, Popp O, Teixeira Alves LG, Reinecke L, Riedel M, Wyler E, Franitza M, Becker K, et al: Rapid UPF1 depletion illuminates the temporal dynamics of the NMD-regulated human transcriptome. Mol Cell 2025, 85:3524–3546 e3512.

5. Raimondeau E, Bufton JC, Schaffitzel C: New insights into the interplay between the translation machinery and nonsense-mediated mRNA decay factors. Biochem Soc Trans 2018, 46:503–512.

6. Parikh VN, Day SM, Lakdawala NK, Adler ED, Olivotto I, Seidman CE, Ho CY: Advances in the study and treatment of genetic cardiomyopathies. Cell 2025, 188:901–918.

7. Lopes LR, Ho CY, Elliott PM: Genetics of hypertrophic cardiomyopathy: established and emerging implications for clinical practice. Eur Heart J 2024, 45:2727–2734.

8. Seeger T, Shrestha R, Lam CK, Chen C, McKeithan WL, Lau E, Wnorowski A, McMullen G, Greenhaw M, Lee J, et al: A Premature Termination Codon Mutation in MYBPC3 Causes Hypertrophic Cardiomyopathy via Chronic Activation of Nonsense-Mediated Decay. Circulation 2019, 139:799–811.

9. Gopalsamy A, Bennett EM, Shi M, Zhang WG, Bard J, Yu K: Identification of pyrimidine derivatives as hSMG-1 inhibitors. Bioorg Med Chem Lett 2012, 22:6636–6641.

10. Li S, Leonard D, Wilkinson MF: T cell receptor (TCR) mini-gene mRNA expression regulated by nonsense codons: a nuclear-associated translation-like mechanism. J Exp Med 1997, 185:985–992.

11. Muhlemann O, Mock-Casagrande CS, Wang J, Li S, Custodio N, Carmo-Fonseca M, Wilkinson MF, Moore MJ: Precursor RNAs harboring nonsense codons accumulate near the site of transcription. Mol Cell 2001, 8:33–43.

12. Wallmeroth D, Lackmann JW, Kueckelmann S, Altmuller J, Dieterich C, Boehm V, Gehring NH: Human UPF3A and UPF3B enable fault-tolerant activation of nonsense-mediated mRNA decay. EMBO J 2022, 41:e109191.

13. Li Y, Wan L, Zhang L, Zhuo Z, Luo X, Cui J, Liu Y, Su F, Tang M, Xiao F: Evaluating the activity of nonsense-mediated RNA decay via Nanopore direct RNA sequencing. Biochem Biophys Res Commun 2022, 621:67–73.

14. Vicente-Crespo M, Palacios IM: Nonsense-mediated mRNA decay and development: shoot the messenger to survive? Biochem Soc Trans 2010, 38:1500–1505.

15. Wittkopp N, Huntzinger E, Weiler C, Sauliere J, Schmidt S, Sonawane M, Izaurralde E: Nonsense-mediated mRNA decay effectors are essential for zebrafish embryonic development and survival. Mol Cell Biol 2009, 29:3517–3528.

16. Medghalchi SM, Frischmeyer PA, Mendell JT, Kelly AG, Lawler AM, Dietz HC: Rent1, a trans-effector of nonsense-mediated mRNA decay, is essential for mammalian embryonic viability. Hum Mol Genet 2001, 10:99–105.

17. Weischenfeldt J, Damgaard I, Bryder D, Theilgaard-Monch K, Thoren LA, Nielsen FC, Jacobsen SE, Nerlov C, Porse BT: NMD is essential for hematopoietic stem and progenitor cells and for eliminating by-products of programmed DNA rearrangements. Genes Dev 2008, 22:1381–1396.

18. Palou-Marquez G, Supek F: Variable efficiency of nonsense-mediated mRNA decay across human tissues, tumors and individuals. Genome Biol 2025, 26:316.

19. Bongiorno R, Colombo MP, Lecis D: Deciphering the nonsense-mediated mRNA decay pathway to identify cancer cell vulnerabilities for effective cancer therapy. J Exp Clin Cancer Res 2021, 40:376.

20. Liu C, Karam R, Zhou Y, Su F, Ji Y, Li G, Xu G, Lu L, Wang C, Song M, et al: The UPF1 RNA surveillance gene is commonly mutated in pancreatic adenosquamous carcinoma. Nat Med 2014, 20:596–598.

21. Bokhari A, Jonchere V, Lagrange A, Bertrand R, Svrcek M, Marisa L, Buhard O, Greene M, Demidova A, Jia J, et al: Targeting nonsense-mediated mRNA decay in colorectal cancers with microsatellite instability. Oncogenesis 2018, 7:70.

22. Burridge PW, Matsa E, Shukla P, Lin ZC, Churko JM, Ebert AD, Lan F, Diecke S, Huber B, Mordwinkin NM, et al: Chemically defined generation of human cardiomyocytes. Nat Methods 2014, 11:855–860.

23. Doroudgar S, Volkers M, Thuerauf DJ, Khan M, Mohsin S, Respress JL, Wang W, Gude N, Muller OJ, Wehrens XH, et al: Hrd1 and ER-Associated Protein Degradation, ERAD, are Critical Elements of the Adaptive ER Stress Response in Cardiac Myocytes. Circ Res 2015, 117:536–546.

24. Mearini G, Stimpel D, Geertz B, Weinberger F, Kramer E, Schlossarek S, Mourot-Filiatre J, Stoehr A, Dutsch A, Wijnker PJ, et al: Mybpc3 gene therapy for neonatal cardiomyopathy enables long-term disease prevention in mice. Nat Commun 2014, 5:5515.

25. Bolger AM, Lohse M, Usadel B: Trimmomatic: a flexible trimmer for Illumina sequence data. Bioinformatics 2014, 30:2114–2120.

26. Dobin A, Davis CA, Schlesinger F, Drenkow J, Zaleski C, Jha S, Batut P, Chaisson M, Gingeras TR: STAR: ultrafast universal RNA-seq aligner. Bioinformatics 2013, 29:15–21.

27. Liao Y, Smyth GK, Shi W: The R package Rsubread is easier, faster, cheaper and better for alignment and quantification of RNA sequencing reads. Nucleic Acids Res 2019, 47:e47.

28. Love MI, Huber W, Anders S: Moderated estimation of fold change and dispersion for RNA-seq data with DESeq2. Genome Biol 2014, 15:550.

29. Lawrence M, Huber W, Pages H, Aboyoun P, Carlson M, Gentleman R, Morgan MT, Carey VJ: Software for computing and annotating genomic ranges. PLoS Comput Biol 2013, 9:e1003118.

30. Anders S, Reyes A, Huber W: Detecting differential usage of exons from RNA-seq data. Genome Res 2012, 22:2008–2017.

31. Reyes A, Anders S, Weatheritt RJ, Gibson TJ, Steinmetz LM, Huber W: Drift and conservation of differential exon usage across tissues in primate species. Proc Natl Acad Sci U S A 2013, 110:15377–15382.

